# Endo-lysosomal Aβ concentration and pH enable formation of Aβ oligomers that potently induce Tau missorting

**DOI:** 10.1101/2020.06.28.175885

**Authors:** Marie P. Schützmann, Filip Hasecke, Sarah Bachmann, Mara Zielinski, Sebastian Hänsch, Gunnar F. Schröder, Hans Zempel, Wolfgang Hoyer

**Author notes:** Correspondence and requests for materials should be addressed to W.H.

## Abstract

Amyloid-β peptide (Aβ) forms metastable oligomers >50 kD, termed AβOs or protofibrils, that are more effective than Aβ amyloid fibrils at triggering Alzheimer’s disease-related processes such as synaptic dysfunction and Tau pathology, including Tau mislocalization. In neurons, Aβ accumulates in endo-lysosomal vesicles at low pH. Here, we show that the rate of AβO assembly is accelerated 8,000-fold upon pH reduction from extracellular to endo-lysosomal pH, at the expense of amyloid fibril formation. The pH-induced promotion of AβO formation and the high endo-lysosomal Aβ concentration together enable extensive AβO formation of Aβ42 under physiological conditions. Exploiting the enhanced AβO formation of the dimeric Aβ variant dimAβ we furthermore demonstrate targeting of AβOs to dendritic spines, potent induction of Tau missorting, a key factor in tauopathies, and impaired neuronal activity. The results suggest that the endosomal/lysosomal system is a major site for the assembly of pathomechanistically relevant AβOs.

## Introduction

Aβ amyloid fibrils are highly stable protein aggregates of regular cross-β structure that constitute the main component of the senile plaques in the brains of Alzheimer’s disease (AD) patients^1–3^. Although amyloid fibrils can exert toxic activities, metastable Aβ oligomers are thought to represent the main toxic species in AD^3–5^. At sufficiently high monomer concentration, Aβ readily forms oligomers with molecular weights >50 kD with spherical, curvilinear, and annular shapes, where the elongated structures appear as “beads-on-a-string”-like assemblies of spherical oligomers^4–11^. While multiple names have been given to these metastable Aβ oligomers, including AβOs, ADDLs, and protofibrils, they seem to be closely related with regard to their structures and detrimental activities, and likely form along a common pathway^6,7,12^. Importantly, this pathway is distinct from that of amyloid fibril formation, i.e., AβOs are not intermediates on the pathway to amyloid fibrils (they are “off-pathway”) but constitute an alternative Aβ assembly type with distinct toxic activities (Fig. 1a) ^4,5,11,13^. The distinct nature of Aβ amyloid fibrils and AβOs is also reflected in their different formation kinetics. Aβ amyloid fibrils form by nucleated polymerization with crucial contributions from secondary nucleation processes, resulting in the characteristic sigmoidal growth time courses that feature an extended lag-time^14^. AβOs, on the other hand, form in a lag-free oligomerization reaction that has a substantially higher monomer concentration dependence than amyloid fibril formation^11^.

**Fig. 1.**
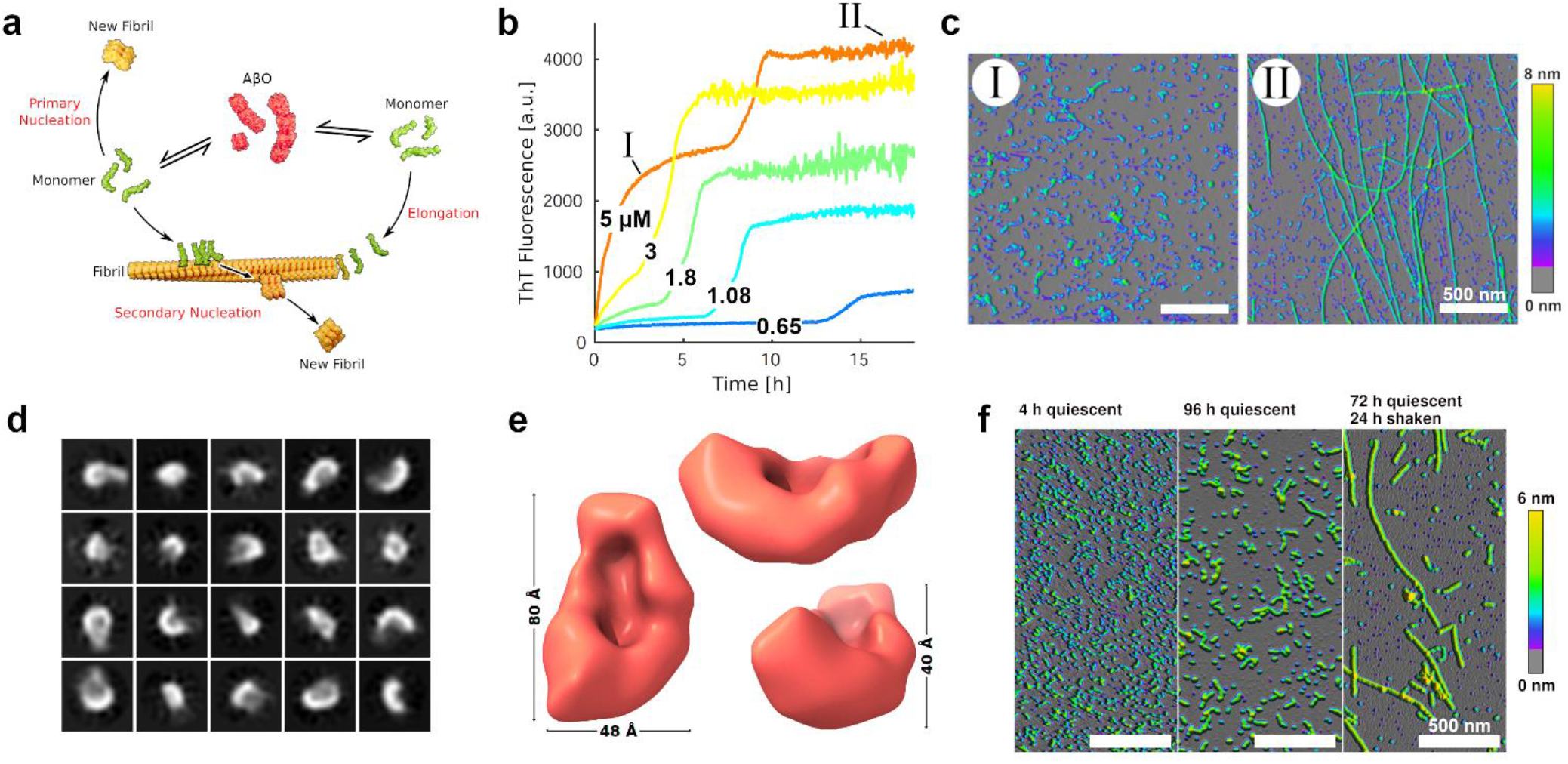
AβOs assemble from dimAβ in a lag-free oligomerization reaction. **a** Scheme of AβO and amyloid fibril formation. **b** Biphasic assembly kinetics of dimAβ at pH 7.4 and indicated concentrations monitored by ThT fluorescence. **c** AFM images corresponding to the two kinetic phases. **d** Exemplary 2D classes of the smallest dimAβ AβO species observed in cryo-EM micrographs. **e** 3D density reconstruction of this dimAβ AβO species at a resolution of 17 Å by cryo-EM. **f** AFM images of dimAβ assemblies formed upon incubation at pH 7.4 in microcentrifuge tubes.

Several lines of evidence support a critical role of AβOs in AD pathogenesis. AβOs of sizes >50 kD are the main soluble Aβ species in biological samples^15^. They are synaptotoxic, disrupt long-term potentiation, and cause cognitive impairment in mouse and non-human primate models^4,8,16–22^. Furthermore, AβOs induce oxidative stress, endoplasmic reticulum stress, neuroinflammation, and elicit Tau missorting, the earliest hallmark of tauopathy in AD ^20,22–28^. The detrimental effects are enhanced by pathogenic Aβ mutations that specifically promote AβO formation, in particular the arctic (Aβ E22G) and the Osaka (Aβ ΔE22) mutations^21,22,27,29,30^. Consequently, targeting AβOs therapeutically is an important alternative to amyloid-centric approaches and has entered clinical evaluation^31–33^.

AβOs were suggested to trigger toxic effects through ligand-like binding to a remarkably high number of candidate receptors^4,34^. AβOs achieve clustering of receptors in cell surface signaling platforms, probably promoted by the multivalency inherent to AβOs^4,34,35^. AβO clustering is especially prominent at dendritic spines, which deteriorate upon prolonged exposure to AβOs^17^. Importantly, this effect is mediated by Tau protein, providing a connection between the Aβ and the Tau aspects of AD pathogenesis. AβOs induce missorting of Tau into the somatodendritic compartment as well as Tau hyperphosphorylation, leading to microtubule destabilization and spine loss^22,36–38^.

In addition to receptor binding of extracellular AβOs, intracellular AβOs are thought to contribute to AD pathogenesis^39^. The endosomal-lysosomal system is the main site not only for Aβ production but also for the uptake of Aβ monomers and AβOs^26,40–47^. Aβ accumulates in endosomes/lysosomes, which promotes aggregation with potential consequences for cellular homeostasis as well as for the spreading of Aβ pathology by exocytosis of aggregated Aβ species^26,27,40,43–45,47–49^.

At neutral pH, high Aβ concentrations are required for AβO formation. Widely-used protocols for AβO preparation start from around 100 μM Aβ monomers^7,8,10^. At tenfold lower Aβ concentration, the formation of AβOs is already greatly disfavored, which enables the investigation of the pure sigmoidal time course of amyloid fibril formation, including the analysis of on-pathway oligomer formation^14,50,51^. These on-pathway oligomers, however, are short-lived, rapidly consumed in the process of fibril formation, and, as evident from the different assembly kinetics, clearly distinct from the neurotoxic off-pathway AβOs introduced above. To investigate AβO formation, we have generated a dimeric variant of Aβ termed dimAβ, in which two Aβ40 units are linked in one polypeptide chain through a flexible glycine-serine-rich linker^11^. In dimAβ, the conformational properties of the Aβ40 units are not altered as compared to free Aβ40 monomers^11^. The linkage of two Aβ units, however, increases the local Aβ concentration, which strongly promotes the highly concentration-dependent formation of AβOs^11^ (Fig. 1b,c). The advantages in applying dimAβ for the study of AβOs are: First, AβOs form already above a threshold concentration (critical oligomer concentration, COC) of ~1.5 μM dimAβ. Second, the increased local Aβ concentration preferentially accelerates AβO formation as compared to Aβ fibril formation, resulting in an enhanced separation of the kinetic phases of AβO and Aβ fibril formation which facilitates analysis.

There is an apparent discrepancy between the obvious pathogenic relevance of AβOs and the high μM Aβ concentrations required for AβO formation at neutral pH *in vitro*, which exceeds the estimated pico-to nanomolar concentrations of extracellular Aβ in normal brain by several orders of magnitude^43^. However, accumulation of Aβ in the endo-lysosomal system was shown to result in micromolar Aβ concentrations in late endosomes and lysosomes^43^, suggesting that these acidic vesicles might be the prime sites of AβO formation. Acidic conditions have been reported to accelerate Aβ aggregation^52^. Here, we applied dimAβ and Aβ42 to test if pH reduction from neutral to endo-lysosomal pH affects AβO formation. We find that endo-lysosomal pH in fact strongly accelerates AβO formation, whereas amyloid fibril formation is delayed, suggesting that AβO formation is the dominant aggregation process in endosomes/lysosomes. We furthermore show that dimAβ is a disease-relevant model construct for pathogenic AβO formation by demonstrating that dimAβ AβOs target dendritic spines, induce AD-like somatodendritic Tau missorting, and reduce synaptic transmission in terminally matured primary neurons. This indicates that dimAβ-derived oligomers are suitable for the study of downstream mechanistic and neuropathological events in the progression of AD.

## Results

### DimAβ assembles into AβOs that bind to dendritic spines and potently induce tau missorting

The assembly kinetics of dimAβ at neutral pH monitored by Thioflavin T (ThT) show a biphasic behavior above a concentration (COC) of ~1.5 μM, with the first phase corresponding to the lag-free oligomerization into AβOs and the second phase reflecting amyloid fibril formation^11^ (Fig. 1b,c). DimAβ AβOs are of spherical and curvilinear shape (Fig. 1c) and rich in β-structure^11^, in agreement with the characteristics of AβOs formed from Aβ40 and Aβ42 (ref. ^4–6,9,13,20^). We applied cryo-EM to further characterize dimAβ AβOs structurally. As larger AβOs seem to be assemblies of small spherical structures, our analysis focused on the smallest AβOs observed in the micrographs (Fig. 1d,e, Supplementary Figs. 1-3). We obtained a 3D density reconstruction (Fig. 1e) at a resolution of 17 Å which shows a bowl-shaped structure with dimensions of 80 x 48 x 40 Å. From this reconstruction we were able to calculate the approximate molecular mass that fits into the density to be 62 kDa (Supplementary Fig. 3; see Methods). Therefore, the smallest AβO species, as visible on the micrographs, likely contains six dimAβ monomers (total MW of 60.2 kD), which corresponds to twelve Aβ40 units. Dodecameric Aβ oligomers were observed before in AβO preparations from synthetic peptide or isolated from AD brain or mouse models, and have been associated with neuronal dysfunction and memory impairment^53–56^.

AβO formation occurred on the same time scale in the plate reader experiment as in microcentrifuge tubes (Fig. 1b,c,f). In contrast, extensive amyloid formation was observed in the plate reader experiment after ~10 h, but was not detectable when AβOs were incubated in microcentrifuge tubes for several days, unless the microcentrifuge tube was agitated (Fig. 1b,c,f). This suggests that the movement of the microplate in the plate reader, caused by scanning of the wells during measurements every 3 minutes and 2 seconds of preceding orbital shaking, creates sufficient agitation to promote amyloid fibril nucleation. When the samples in the microplate were covered with a layer of mineral oil, AβO formation was unaffected but amyloid fibril formation was completely abrogated (Supplementary Fig. 4), in line with the essential role of the air-water interface in Aβ amyloid formation *in vitro*^57^. The strong effects of agitation^14^ and air-water interface on Aβ amyloid fibril formation but not on AβO formation confirms again that their assembly mechanisms are different, and is in line with the notion that AβO formation does not involve a nucleation step^11,58^. When AβOs, formed by incubation of dimAβ above the COC, were diluted to sub-COC concentrations, they persisted for >24 hours, indicating high kinetic stability (Supplementary Fig. 5). We conclude that AβOs formed from dimAβ under quiescent conditions are kinetically stable, not replaced by amyloid fibrils for several days, and can be applied at sub-μM concentrations. DimAβ AβOs may therefore serve as a favorable AβO model.

To test if dimAβ AβOs cause the same biological effects as reported for AβOs formed from Aβ40 or Aβ42, we investigated their binding to dendritic spines, their direct cytotoxicity, their capacity to induce Tau missorting, and their consequences for neuronal function. AβOs were formed from 20 μM dimAβ and added to primary mouse neurons (DIV15-22) to a final concentration of 0.5 μM (all dimAβ AβO concentrations given in dimAβ equivalents). 1 μM Aβ40 was used as monomeric control. DimAβ localized to neuronal dendrites both after 3 and 24 hours of treatment, where it partially co-localized with dendritic protrusions positive for filamentous actin (stained by phalloidin), which mark synaptic spines (Fig. 2a). In contrast, Aβ40 monomers did not show substantial localization to dendrites (Fig. 2a). Direct cytotoxicity was tested by analysis of the sizes and shapes of neuronal nuclei upon staining with NucBlue. The fractions of normal and dense nuclei did not change significantly after incubation with dimAβ AβOs (Fig. 2b), indicating the absence of direct cytotoxicity, in line with previous reports on AβOs^59^.

**Fig. 2.**
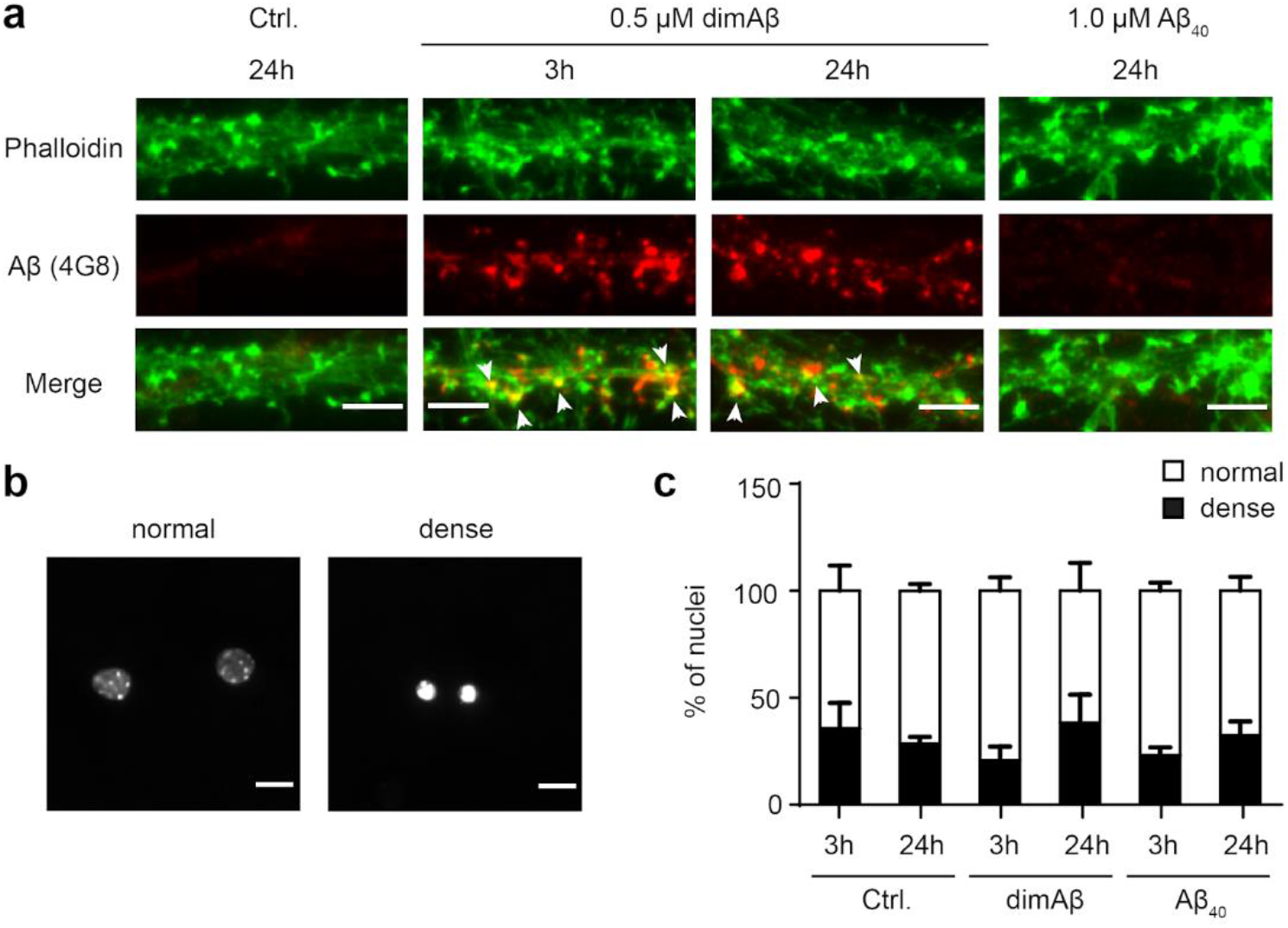
DimAβ AβOs bind to dendrites and postsynaptic spines but has no direct cytotoxic effect on primary mouse neurons. Primary mouse neurons (DIV15-22) were treated with 0.5 μM dimAβ AβOs or 1 μM Aβ40 for 3 and 24 hours. **a** DimAβ AβOs localized to neuronal dendrites both after 3 and 24 hours of treatment, where they partially co-localized with phalloidin, a marker for synaptic spines. Arrows indicate co-localization of dimAβ with phalloidin. Scale bar, 5 μm. **b** Nuclei of primary neurons were stained with NucBlue and analyzed with respect to shape and size. Representative images of normal and dense nuclei. Scale bar, 10 μm. **c** Quantification of normal and dense nuclei of primary neurons after control, Aβ40, or dimAβ AβO treatment revealed no direct cytotoxicity. Around 300 nuclei were analyzed for each condition. Error bars represent SEM. Statistical analysis was done by two-way ANOVA with Tukey’s test for multiple comparisons.

Tau cellular distribution was analyzed with an anti-Tau (K9JA) antibody. DimAβ treated neurons showed strong enhancement of the fluorescence signal of Tau in the soma after 24 hours of treatment (Fig. 3), indicating pathological somatodendritic Tau missorting as previously reported for AβOs^37,38^. In contrast, Aβ40 monomers did not induce Tau missorting in our experimental setting (Fig. 3). In previous studies, Tau missorting and spine loss were reversible within 12-24 hours due to loss of AβO potency (transformation of AβOs over time to larger, non-toxic aggregates)^37,60^. Here, we observe an increase of Tau missorting over time, which indicates remarkable kinetic stability and persistent activity in terms of capacity to induce AD-like Tau missorting of dimAβ AβOs.

**Fig. 3.**
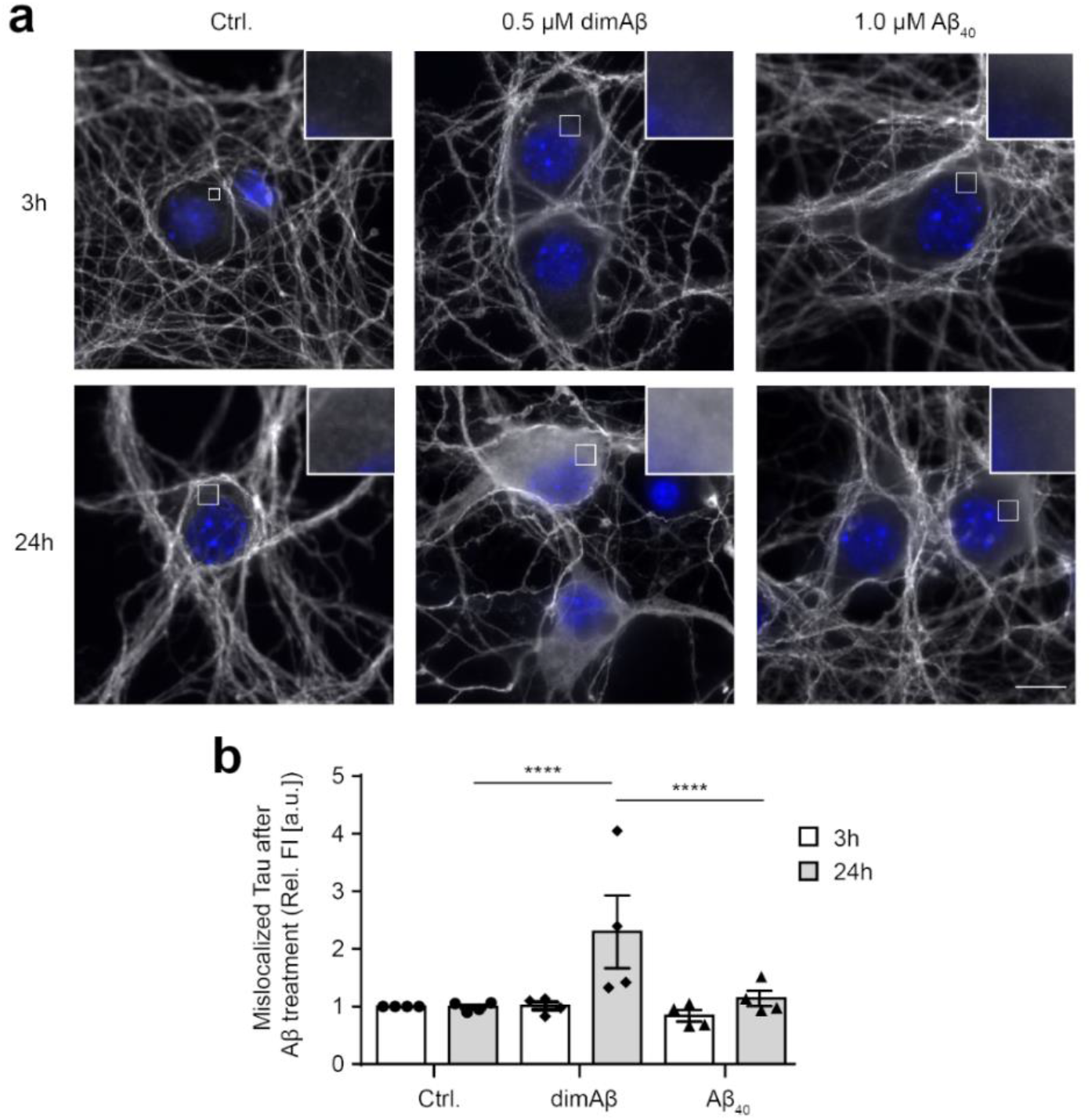
DimAβ AβOs induce pathological somatodendritic missorting of Tau. Primary mouse neurons (DIV15-22) were treated with 0.5 μM dimAβ AβOs or 1 μM Aβ40 for 3 and 24 hours. **a** Representative images of cell bodies of primary neurons after treatment with Aβ. Neurons were stained with anti-Tau (K9JA) antibody, nuclei were stained with NucBlue. DimAβ AβO treated neurons show strong enrichment of fluorescence signal of Tau in the soma only after 24 hours of treatment. Insets show magnification of white boxed areas in the somata. Scalebar, 10 μm. **b** Quantification of Tau enrichment in the soma of primary neurons. Fluorescence intensities of cell bodies were quantified and normalized to control treated neurons after 3 hours of treatment. N=4, 30 cells were analyzed for each condition. Error bars represent SEM. Statistical analysis was done by two-way ANOVA with Tukey’s test for multiple comparisons. Statistical significance: **** = p ≤ 0.0001.

Next, we investigated the consequences of AβO exposure for neuronal function. As readout, we measured spontaneous calcium oscillations in our neuronal cultures after AβO treatment as an indicator for neuronal activity with live-imaging, using the fluorescent cell permeable calcium indicator Fluo4 as previously described^37^. A significant decrease of calcium oscillations was observed after 24h, but not after 3h, of treatment with dimAβ AβOs (Fig. 4). As calcium oscillations in our conditions depend on action potentials and neurotransmission, this indicates that dimAβ AβOs impair neuronal function. With regard to dendritic spine binding, lack of direct cytotoxicity, potent induction of Tau missorting as well as decreased neuronal activity, dimAβ AβOs thus faithfully reproduce the observations made for AβOs formed from Aβ40 or Aβ42, or from 7:3 Aβ40:Aβ42 mixtures regarded as particularly toxic^37^. Of note, dimAβ AβOs effects appeared later (24h vs. 3h) than for the previously studied oligomers, hinting towards their kinetic and structural stability in cell culture conditions.

**Fig. 4.**
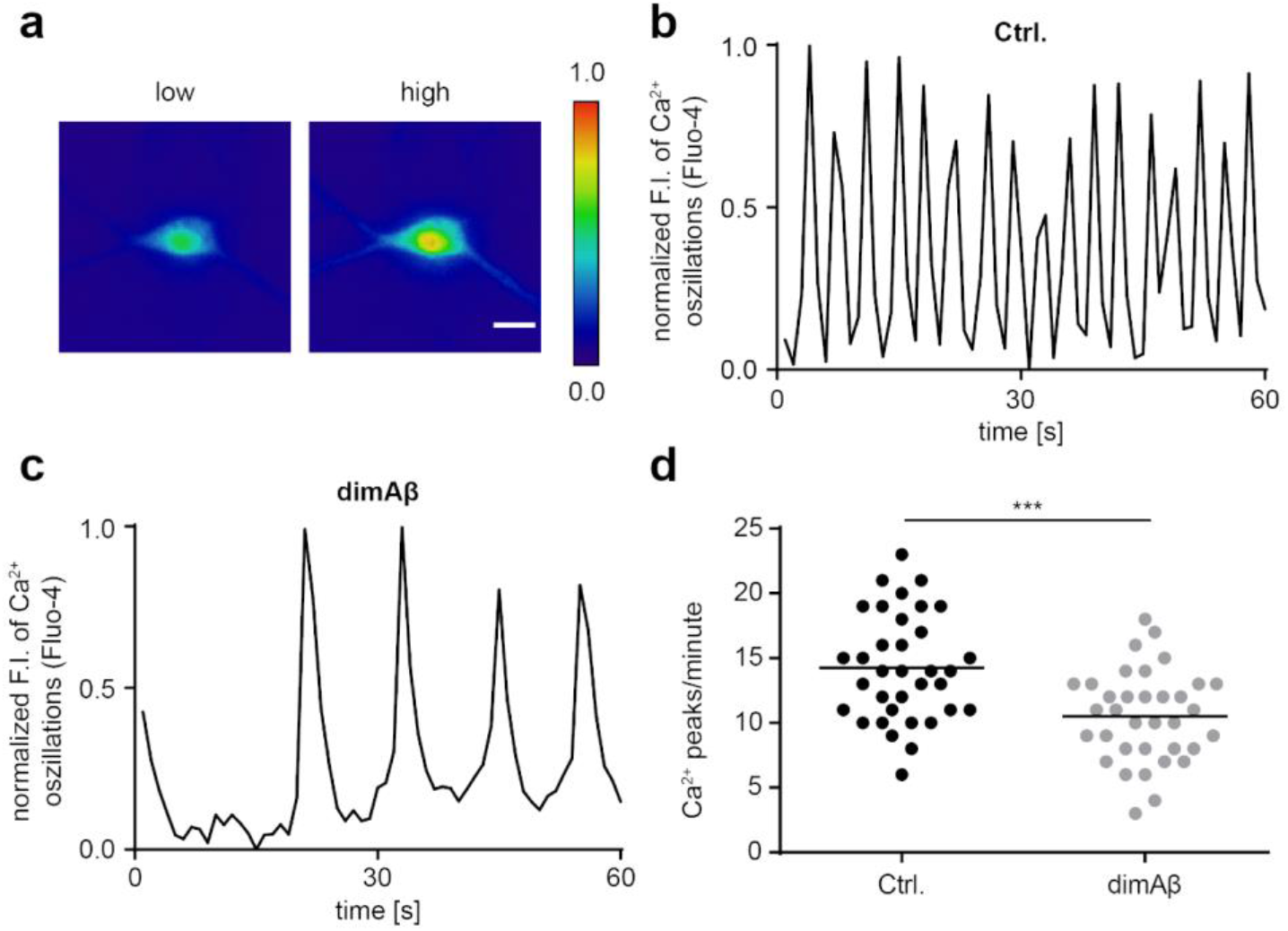
DimAβ AβOs decrease spontaneous calcium oscillations of primary mouse neurons. Primary mouse neurons (DIV15-22) were treated with 0.5 μM dimAβ AβOs for 24 hours. Cells were labelled with calcium-sensitive Fluo-4 dye and spontaneous calcium oscillations were recorded by time-lapse movies. **a** Representative ratiometric images of low and high calcium concentrations in the soma of a neuron. Scalebar, 20 μm. **b**, **c** Representative graphs of spontaneous Ca^2+^ oscillations in **b** vehicle control and **c** dimAβ AβO treated primary neurons. Fluorescence intensities were normalized to minimum values and plotted over time. **d** Quantification of spontaneous Ca^2+^ oscillations in primary neurons after vehicle control or dimAβ AβO treatment. Fluorescence intensities were normalized to minimum values and peaks per minute were counted for each sample. In total, 35 cells were analyzed; statistical analysis was done by unpaired t-test. Statistical significance: *** = p ≤ 0.001.

### Aβ42 monomers as well as dimAβ AβOs accumulate within endo-lysosomal compartments

Next, we aimed to test the uptake of dimAβ AβOs in neuronal cells. First, SH-SY5Y neuroblastoma cells were subjected to 100 nM HiLyte Fluor 647-labeled Aβ42. After 24 hours of incubation Aβ42 accumulated within vesicular foci within the cytoplasm of the cells. Co-staining with a LysoTracker dye showed prominent colocalization suggesting the accumulation of Aβ42 within endo-lysosomal compartments (Fig. 5a). This was in line with findings of previous studies that showed Aβ42 accumulation in acidic vesicles of neuroblastoma cells and primary murine cortical neurons^40,43–45^. Hu et al. measured local Aβ42 concentrations higher than 2.5 μM within endo-lysosomal compartments which exceeds the extracellular concentration by approximately four orders of magnitude^43^.

**Fig. 5.**
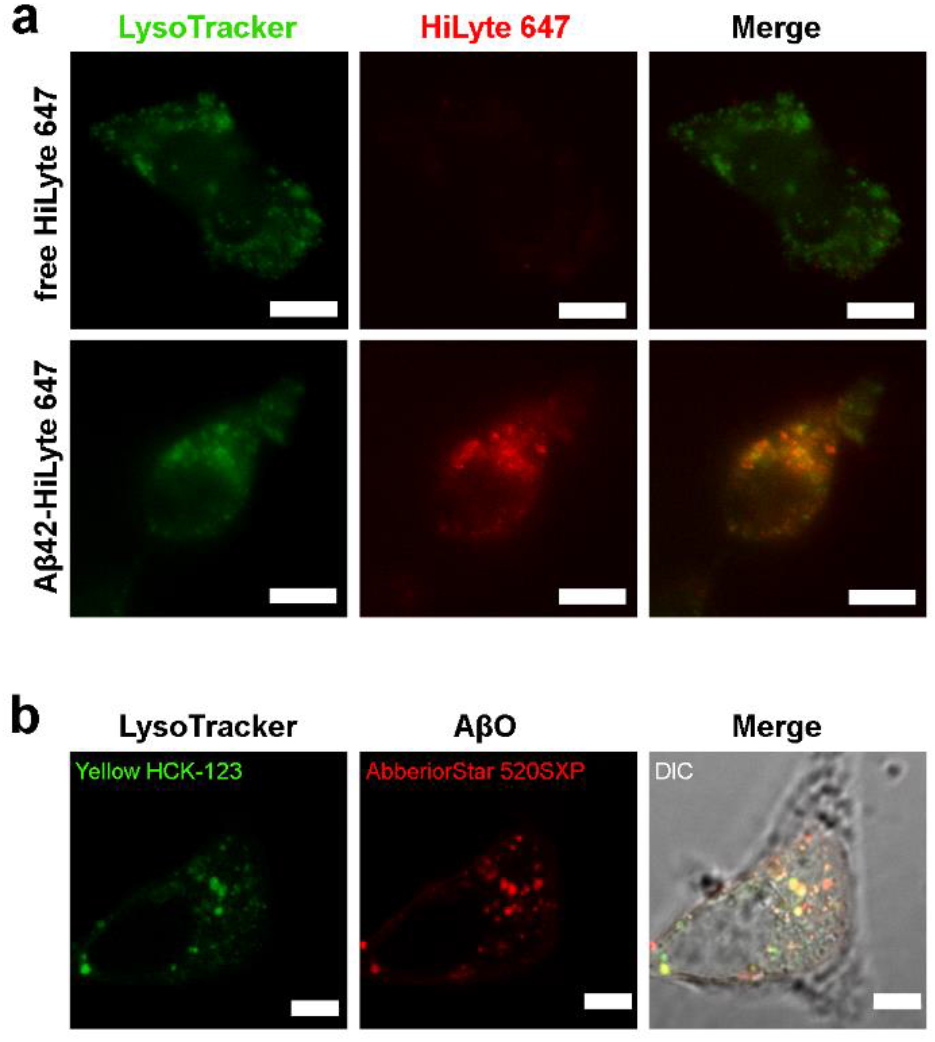
Aβ42 and dimAβ AβOs accumulate in endosomes/lysosomes. SH-SY5Y cells were treated with Aβ42 or dimAβ and co-localization with endo-lysosomal compartments was analyzed. **a** 0.1 μM free HiLyte 647 dye or 0.1 μM HiLyte 647-labeled Aβ42 was added to the cells. After 24 hours the medium was exchanged with fresh medium supplemented with 50 nM Yellow HCK-123 LysoTracker dye. Scalebar, 10 μm. **b** Cells were treated with dimAβ AβOs (0.1 μM AbberiorStar 520SXP-labeled dimAβ + 1 μM unlabeled dimAβ). After 24 hours the medium was exchanged with fresh medium supplemented with 50 nM Yellow HCK-123 LysoTracker dye. Scalebar, 5 μm.

In a second attempt, SH-SY5Y cells were treated with Abberior Star 520SXP-labelled dimAβ AβOs revealing a similar colocalization in acidic vesicles (Fig. 5b). This confirms that both Aβ monomers and AβOs are readily taken up by neuron-like cells and accumulate in the endo-lysosomal system.

### Endo-lysosomal pH promotes AβO assembly but delays amyloid fibril formation

Due to the accumulation of Aβ, endosomes/lysosomes might constitute the dominant site of the highly concentration-dependent AβO formation. Apart from the increased Aβ concentration in endosomes/lysosomes, the low pH in late endosomes (~5.5) and lysosomes (~4.5) might promote AβO formation. We used dimAβ to simultaneously determine the specific effects of pH on AβO formation and on amyloid fibril formation. Lyophilized dimAβ was dissolved in 6 M buffered guanidinium chloride, followed by size-exclusion chromatography (SEC) into 1 mM NaOH, leading to a pH of 10.9, and added to the wells of a microplate. The basic pH conditions prohibit premature aggregation of Aβ^61^. The pH-dependent aggregation reaction was initiated in the microplate reader by injection of a 10x buffer yielding the desired final pH, allowing for monitoring of ThT fluorescence without any substantial delay. We determined the kinetics of dimAβ assembly between pH 4.8 and 7.6 in the concentration range 0.65 – 5.0 μM. At neutral pH, the initial kinetic phase reflecting AβO formation spanned several hours, but upon pH reduction AβO formation was continuously accelerated and occurred within a few seconds at pH 4.8 (Fig. 6a-g). ThT fluorescence intensity decreased at acidic pH^62^, but was still sufficiently sensitive to detect the signal of AβO formation at pH 4.8 and 0.65 μM dimAβ (Fig. 6g). For pH 7.4, we have previously shown that a global fit of an *n*^th^-order oligomerization reaction to the concentration-dependent assembly kinetics is in good agreement with the data and yields a reaction order of ~3.3 for dimAβ AβO formation^11^. Here, we found that a reaction order of 3 applied to global fitting of the concentration-dependent data results in fits that reproduce the kinetic traces at all pH values (Fig. 6a-g). This indicates that the fundamental mechanism of AβO formation is not affected by pH reduction. A logarithmic plot of the obtained oligomerization rate constants against pH shows a linear trend with a slope of −1.56, i.e., the rate constant decreases 36-fold per pH unit within the investigated pH range (Fig. 6h). At pH 4.8, in between lysosomal and endosomal pH, AβO formation is 7,900-fold faster than at interstitial pH (7.3).

**Fig. 6.**
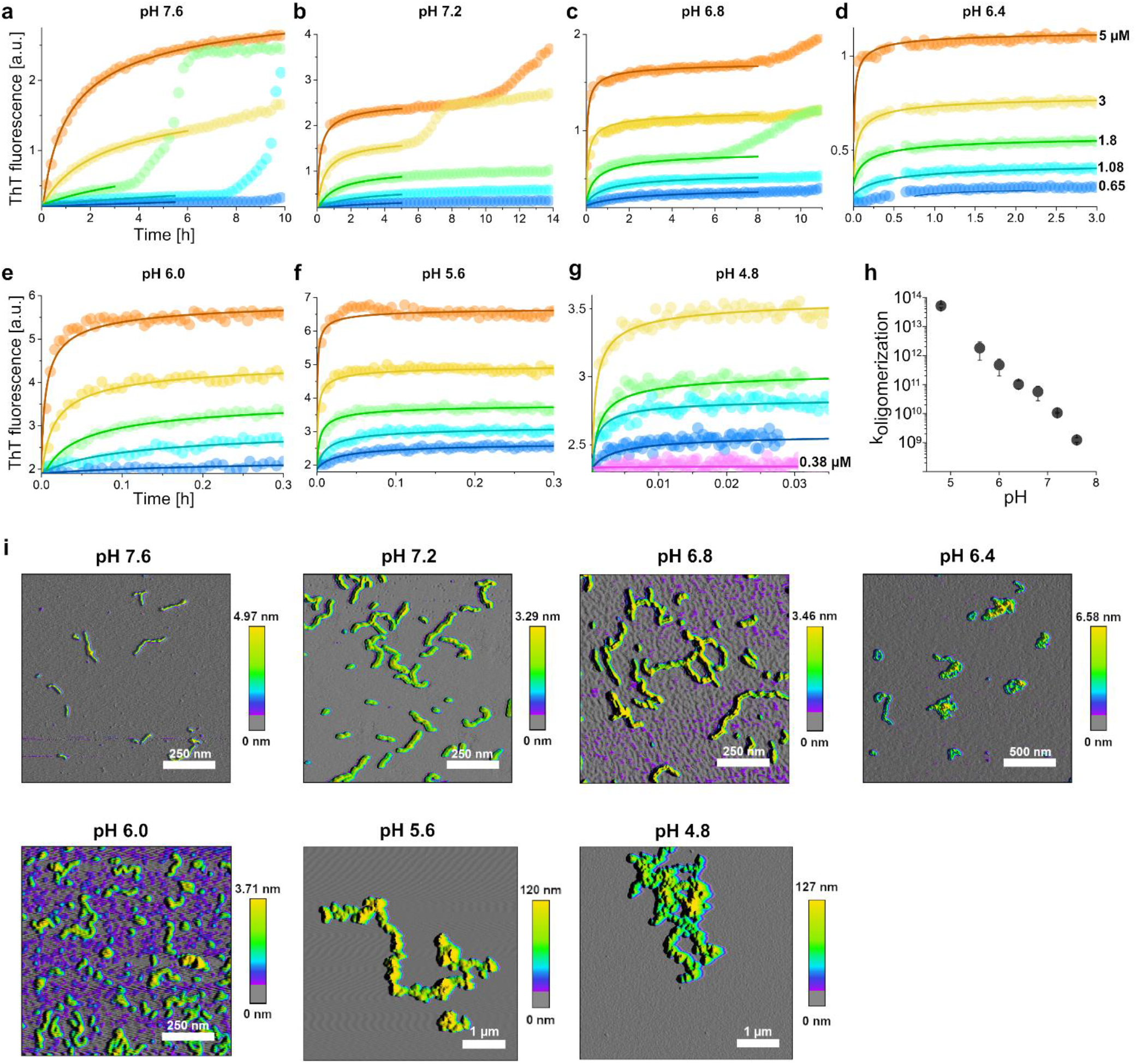
pH dependence of dimAβ assembly kinetics. **a-g** DimAβ assembly at concentrations between 0.65 and 5 μM and at pH values between 4.8 and 7.6 monitored by ThT fluorescence. Solid lines represent global fits to the data using a one-step oligomerization model with a shared reaction order of 3 for all pH values and concentrations, and an individual oligomerization rate constant per pH value. **h** Logarithmic plot of the obtained oligomerization rate constants vs. pH. **i** AFM images of dimAβ AβO formed at the different pH values. Note the dramatic change in the height scale bar upon pH decrease to <6.0 due to formation of large AβO clusters.

AβOs formed at different pH values were imaged by AFM (Fig. 6i). From pH 7.6 to pH 6.8 AβOs were mainly spherical and curvilinear structures, the latter apparently resulting from bead-chain-like association of the spherical AβOs^6^. At pH 6.4, AβOs showed an increased tendency to form more compact structures, such as annular protofibrils and denser clusters. Below pH 6.0, AβOs associated into large clusters, in line with a previous description of Aβ40 aggregates at pH 5.8^52^. Thus, while the fundamental mechanism of AβO formation seems to be unaffected by pH reduction, there is an additional level of particle aggregation involved below pH 6.0.

The second kinetic phase in the ThT time course of dimAβ aggregation reports on amyloid fibril formation^11^. It is characterized by a lag-time, which reflects the primary and secondary nucleation events involved in nucleated polymerization^14,50^. In contrast to the acceleration of AβO formation, the lag-time of amyloid formation did not decrease with decreasing pH. On the contrary, the amyloid fibril formation phase could not be observed within 10 hour experiments at pH values of 6.8 and below. This can be explained by the inhibition that the rapidly forming AβOs entail on amyloid formation: First, AβOs compete for the monomer growth substrate of amyloid fibril growth; second, AβOs actively inhibit amyloid fibril growth^11^.

### AβO assembly of Aβ42 is enabled under endo-lysosomal conditions

We investigated if the promotion of AβO formation at endo-lysosomal pH is sufficient to also support AβO formation from Aβ42 at relevant endo-lysosomal Aβ concentrations, determined to be well above 2.5 μM^43^. At pH 7.2, Aβ42 in the concentration range 1.9-9 μM displayed sigmoidal assembly kinetics typical for amyloid fibril formation (Fig. 7a). In contrast, at pH 4.5 lag-free aggregation occurred at a concentration of 5.4 μM and above (Fig. 7b). The change from lag-containing to lag-free conditions at pH 4.5 was accompanied by a switch in aggregate morphology from amyloid fibril networks to large AβO clusters identical to those observed for dimAβ at endo-lysosomal pH (Fig. 7c,d). This indicates that under endo-lysosomal conditions the local Aβ concentration can exceed the COC of AβO formation, suggesting that endosomes/lysosomes may represent crucial sites of AβO formation *in vivo*.

**Fig. 7.**
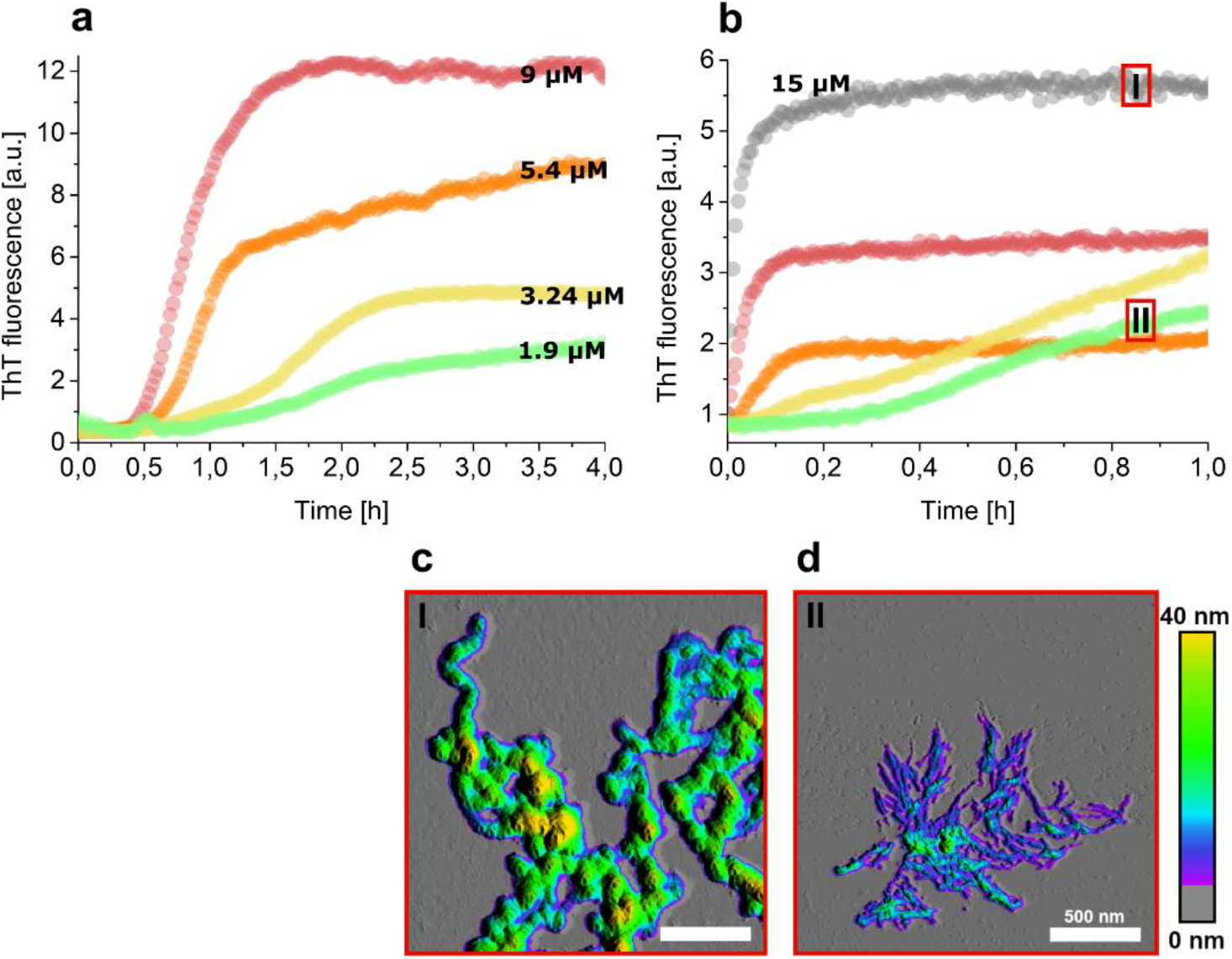
Aβ42 rapidly forms AβOs at endo-lysosomal pH. **a**, **b** Aβ42 assembly at **a** pH 7.2 or **b** pH 4.5 at concentrations between 1.9 and 15 μM monitored by ThT fluorescence. **c**, **d** AFM images of **c** AβOs formed of 15 μM Aβ42 and **d** amyloid fibril networks formed of 1.9 μM Aβ42, both at pH 4.5.

Aβ aggregates can leak from endosomes/lysosomes into the cytosol and to other cell compartments or can be secreted and spread to other cells, potentially contributing to the propagation of Aβ pathology^26,27,43,44,49^. Upon transfer from endosomes/lysosomes to the cytosol or interstitial fluid, AβOs experience a shift from acidic to neutral pH. We tested the kinetic stability of AβOs formed at pH 4.5 after a shift to neutral pH by monitoring the ThT intensity and by imaging the aggregate morphology by AFM. We applied Aβ42 at a concentration of 10 μM in this experiment, as Aβ42 does not form AβOs *de novo* at this concentration at neutral pH. Any AβOs observed after the pH shift can therefore safely be ascribed to the kinetic stability of AβOs pre-formed under acidic conditions. As before, a pH shift from basic pH to pH 4.5 was applied to initiate AβO formation. After AβO formation had reached a steady state, pH was adjusted to 7.2 by a further injection of a corresponding buffer stock. After the adjustment to neutral pH, there was an instantaneous increase in ThT fluorescence (Supplementary Fig. 6), which can be explained by the pH dependence of ThT fluorescence^62^. Thereafter, the ThT fluorescence did not exhibit any other larger changes that would be expected in the case of disassembly of AβOs or replacement of AβOs by an alternative type of aggregate. Apart from dense clusters like those observed for low pH AβOs, AFM images showed spherical and curvilinear structures typical for AβOs formed at neutral pH, indicating dissociation of the AβO clusters into their constituents (Fig. 8a). In fact, the AFM images suggest that smaller AβOs detach from fraying AβO clusters. Taken together, the ThT and AFM data demonstrate that AβOs formed at endo-lysosomal pH possess a high kinetic stability after shifting to neutral pH, which is, however, accompanied by dissociation of large AβO clusters into spherical and curvilinear AβOs.

**Fig. 8.**
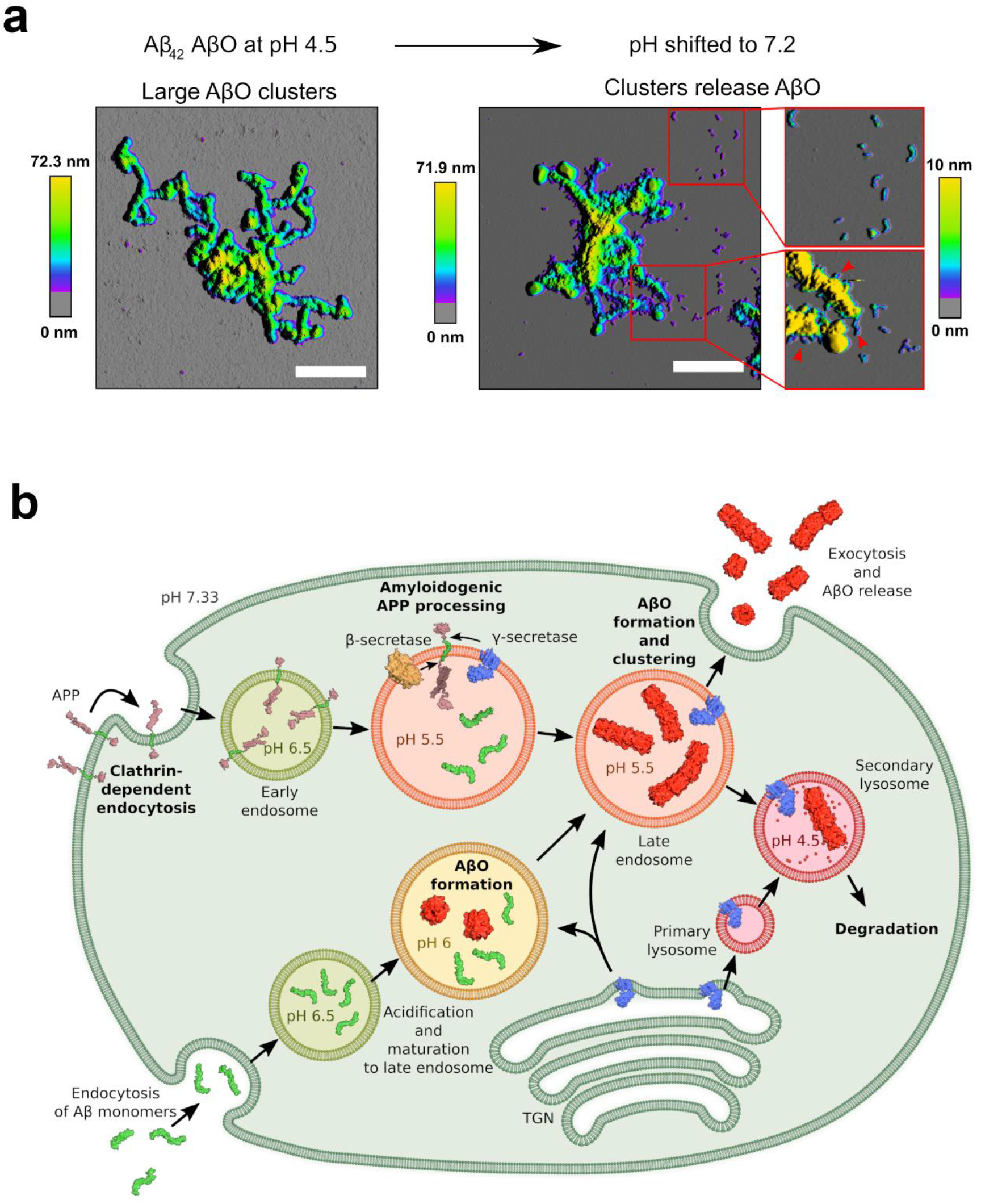
Stability of AβOs formed from Aβ42 at endo-lysosomal pH after shifting to neutral pH. **a** AFM images of AβOs formed of 10 μM Aβ42 at pH 4.5 before (left) and after (right) shift to pH 7.2. Red arrowheads point to a few of the sites where AβOs seem to detach from AβO clusters. Scalebar, 500 nm. **b** Scheme of intracellular APP processing and Aβ uptake^47,72,73^, including potential formation of AβOs especially in endo-lysosomal compartments. Protein structure images were prepared using pdb entries 1OWT, 1IYT, 1RW6, 3DXC, 4UIS, 1SGZ.

## Discussion

AβOs have been identified as the main neurotoxic Aβ species in AD. The characterization of the most critical disease-related AβOs has revealed that they are metastable oligomers >50 kD in size that do not represent intermediates of amyloid fibril formation but are an alternative Aβ assembly type. However, the conditions required for AβO formation and the underlying mechanism have not been elucidated in detail. Here, we show that AβO formation is highly pH-dependent, and is accelerated ~8,000-fold upon a change in pH from neutral to endo-lysosomal pH. This enables AβO formation at physiologically relevant Aβ concentrations. The strong acceleration of AβO formation at pH 4.5-5.5 suggests that the endosomal/lysosomal system might be a major site of AβO formation. AβOs may either form from Aβ monomers that have been newly generated by APP processing or from endocytosed monomers (Fig. 8b)^39–41,43,46,47^. APP processing in endo-lysosomal compartments by γ-secretase containing presenilin 2 generates a prominent pool of intracellular Aβ that is enriched in Aβ42 (ref. ^47^). Esbjörner et al. applied fluorescence lifetime and superresolution imaging to determine the kinetics of Aβ aggregation in live cells and found that aggregation occurred in endo-lysosomal compartments^40^. Importantly, they reported that Aβ42 aggregated without a lag-time into compact, dense structures^40^. Both the absence of a lag-time and the structural characterization are in line with the low pH AβO clusters described here, suggesting that AβO clusters indeed form in endo-lysosomal compartments and represent the dominant Aβ aggregate species in live cells. Subsequently, AβOs might cause lysosomal impairment, leak into the cytosol and cause intracellular damage, or might be secreted and spread to neighboring cells, where they could contribute to the propagation of pathology^39,41,43–45,47^.

Enhanced aggregation at acidic pH is a known property of Aβ with established relevance for sample preparation^61^. Our results are in line with a study on the aggregation of Aβ40 (at a concentration of 230 μM) at pH 5.8 that reported the rapid formation of large clusters with (proto)fibrillar and globular substructures that were not able to seed, but rather inhibited, amyloid fibril formation^52^. Our analysis of the aggregation kinetics reveals that these low pH Aβ aggregates, often termed amorphous aggregates, form along the same pathway as neutral pH AβOs and therefore represent particle aggregates of AβOs. This is supported by the observation that low pH AβO clusters release spherical and curvilinear AβOs upon a shift to neutral pH (Fig. 8a).

The increasing clustering of AβOs upon pH reduction from neutral to pH 6 points to the high propensity of AβOs to associate. At neutral pH, self-association of spherical AβOs results in curvilinear assemblies. A decrease of pH leads to an increase in annular and compact assemblies and finally to large AβO clusters (Fig. 6). This propensity of AβOs to associate likely also contributes to their clustering with neuronal receptors^34,35^ and to their accumulation around amyloid fibril plaques^63^.

In contrast to AβO formation, amyloid fibril formation of dimAβ is slowed down at acidic pH. This pH dependence is not an inherent property of Aβ amyloid fibril formation: In the absence of AβOs, Aβ42 amyloid fibril formation occurs rapidly at pH 4.5 (Fig. 7b, 1.9 μM trace). Delayed amyloid fibril formation upon pH reduction is only observed in combination with accelerated AβO formation, and can be explained by the two inhibitory activities of AβOs on amyloid fibril formation: AβOs compete with amyloid fibrils for monomers (Fig. 1a) and furthermore inhibit amyloid fibril growth actively^11^.

DimAβ AβOs show dendritic spine binding, lack direct cytotoxicity, potently induce Tau missorting, and decrease neuronal activity, suggesting that they constitute a suitable AβO model construct to study the pathomechanism of AD. Previous AβO preparations showed a loss of potency to induce Tau missorting within 12 hours due to transformation to non-toxic larger Aβ aggregates^37,60^. In contrast, dimAβ AβOs led to extensive and persistent Tau missorting 24 h after application. The sustained activity of dimAβ AβOs is likely a consequence of the kinetic stabilization of the AβO state achieved by the dimer linkage. DimAβ might therefore be an advantageous model for eliciting Tau missorting and downstream consequences, as it represents a model of chronic stress corresponding to the human disease rather than acute insult.

## Methods

### Preparation of dimAβ

DimAβ was recombinantly produced as previously described^11^. For aggregation kinetics experiments, the lyophilized protein was reconstituted in 6M guanidinium chloride, 50 mM sodium-phosphate buffer, pH 7.4, and incubated at room temperature for 30 min. Subsequently, SEC was performed using a Superdex 75 increase column (GE Healthcare) equilibrated with 1 mM NaOH. The concentration of the monomeric dimAβ in the alkaline eluate was measured via tyrosine fluorescence using a pH-adjusted extinction coefficient of 2685 M^−1^ cm^−1^. Samples were always kept on ice until further needed.

### ThT aggregation kinetics

ThT, NaN_3_, NaCl, protein, and 1 mM NaOH were given into the wells of a 96-well low-binding plate (Greiner) such that if filled up to 100 μL, concentrations of 1 μM ThT, 0.02% NaN_3_, 150 mM NaCl, and the desired final protein concentration were reached. The outermost wells of the plate were left blank due to the risk of aberrant aggregation behavior. The plate was put in a BMG ClarioStar platereader fitted with two injectors and tempered at 37°C. One syringe of the injector was equilibrated with 1 mL 10x buffer concentrate. The reaction was started using the injector of the platereader by dispensing 10 μL of the concentrate at highest available speed into each of the wells. This adjusted the pH value *in situ* and initiated oligomerization. Data points were measured in evenly spaced intervals depending on the velocity of the reaction. For shifting the pH *in situ* twice, both syringes were equilibrated with 10x buffer concentrate; the first one resulting in a final buffer concentration of 20 mM and pH 4.5 and the second one resulting in a final buffer concentration of 50 mM and pH 7.2. The first syringe was used to inject 10 μl to initiate oligomerization, whereas the second one was used to inject 11 μl to achieve the shift to neutral pH at a time point were the oligomerization reaction had reached its plateau.

For analysis of the kinetics of AβO formation, the initial phase of the ThT kinetics was fit to one-step oligomerization *n* M → M_n_ (ref. ^11^). The AβO mass concentration, *M_AβO_*, evolves in time according to the following expression

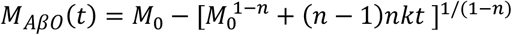

with *M*_0_ the total protein concentration, *k* the oligomerization rate constant, and *n* the oligomer size or reaction order. Global fits to the pH- and concentration-dependent AβO formation data were performed with a reaction order of *n*=3 shared between all data sets, and the oligomerization rate constant *k* as a pH-dependent parameter which was shared within the concentration dependency data sets at a given pH. The proportionality constant relating *M*(t) to ThT fluorescence intensity was treated as a fit parameter with an individual value for every sample.

### Atomic force microscopy (AFM)

10 μl of the dimAβ samples were taken directly from the plate after the ThT assays at a concentration of 5 μM and applied onto freshly cleaved muscovite mica. They were left to dry, washed with 500 μl ddH_2_O, and dried with a stream of N_2_ gas. For imaging dimAβ at pH 4.8, the aforementioned method did not work, likely due to sticking of the sample to the well. Instead, all reaction components apart from the buffer concentrate were premixed and loaded into a micro pipette tip. By adding the reaction components to a vial containing the buffer concentrate and thorough mixing, the reaction was started, before pulling the solution back into the tip. Immediately afterwards, the micropipette was relocated into a 37°C incubation cabinet, where a drop was pushed out to the point where it still stuck to the tip. After 45 sec, the drop was pushed onto the freshly cleaved muscovite mica and preparation commenced as with the other pH values.

For the Aβ42 samples, 5 μl of the respective concentrations were taken, applied onto freshly cleaved muscovite mica and left to dry for 15 min before carefully washing with 200 μl ddH_2_O and drying under a stream of N_2_ gas.

Imaging was performed in intermittent contact mode (AC mode) in a JPK Nano Wizard 3 atomic force microscope (JPK, Berlin) using a silicon cantilever with silicon tip (OMCL-AC160TS-R3, Olympus) with a typical tip radius of 9 ± 2 nm, a force constant of 26 N/m and resonance frequency around 250 kHz. The images were processed using JPK DP Data Processing Software (version spm-5.0.84). For the presented height profiles a polynomial fit was subtracted from each scan line first independently and then using limited data range. False-color height images were overlaid onto the amplitude profile.

### Cryo-EM

For cryo-EM imaging, the AβO sample was plunge-frozen on glow-discharged Quantifoil 1.2/1.3 grids. In total 1308 micrographs were recorded as focal pairs at high defocus (6 μm) and low defocus (using a range of −0.5 to −2 μm) on a Tecnai Arctica (200 kV) using a Falcon III direct electron detector, yielding a pixel size of 0.935 Å. Particle selection was performed automatically using crYOLO^64^. In total 32,211 particles were selected on the high defocus micrographs. The contrast transfer function (CTF) of the micrographs was determined using CTFFIND4^65^. Further image processing was performed using the software package RELION 3.0.5^66^. 2D and 3D classification was conducted on the high defocus images to clean the dataset. A box size of 128 pix, which corresponds to 119.7 Å, and a radial mask with a diameter of 100 Å were used.

The high-defocus micrographs were aligned to the low-defocus micrographs. The relative shifts obtained from this alignment were applied to all particles (that were picked from the high-defocus micrographs) and then the particles were extracted from the low-defocus micrographs with the shifted particle coordinates, while keeping the Euler angles from the high-defocus 3D refinements. A 3D reconstruction calculated from the high defocus images was low-pass filtered to 60 Å and was used as an initial model for further low-defocus 3D refinements. For further processing steps only micrographs that contain a signal beyond a resolution of 5 Å were used. The final resolution of 17 Å was assessed by Fourier shell correlation.

In order to obtain an estimate for the molecular mass within the reconstructed density, 110 pseudo-atomic models with varying number of pseudo-atoms (molecular masses between 10 and 120 kDa) were generated from the density map using the program VISDEM^67^, which is part of the software package DireX^68^. In VISDEM, atoms are randomly placed into a density region with density above a provided threshold. The density threshold was set to yield a volume such that the mass density is fixed at 0.714 ml/g (average mass density observed in proteins). The pseudo-atomic model has a composition of 62.2% C atoms, 20.6% O atoms and 17.2% N atoms, which corresponds to the average composition observed in proteins. Afterwards, a density map was computed from every of the 110 pseudo-atomic models. The VISDEM method was used to sharpen these pseudo-atomic model-maps as well as the EM reconstruction. The sharpening was performed with a resolution cutoff of 17 Å and the mass of the corresponding pseudo-atomic model. Finally, the cross-correlation between the sharpened EM reconstruction and the sharpened pseudo-atomic model-map was computed and plotted for each tested mass. The highest cross-correlation was found for the pseudo-atomic model-map that contains a molecular mass of 62 kDa. One dimAβ monomer (101 amino acids) has a molecular mass of 10.0 kDa. Thus, the reconstructed density likely holds 6 dimAβ monomers. The final 3D reconstruction of the oligomer was sharpened by VISDEM using a mass of 62 kDa and a resolution cutoff of 17 Å.

### Aβ preparations and treatment of primary neurons

Aβ preparations were performed under sterile conditions. DimAβ lyophilisate was resuspended in 50 mM NaOH until completely dissolved. Next, PBS and 50 mM HCl were added and immediately mixed, obtaining a final concentration of 20 μM dimAβ and 40 μM Aβ40, respectively. To induce AβO formation, dimAβ was incubated at 37°C for 16 hours. Aβ40 controls were prepared in the same manner without subsequent incubation. Primary neurons (DIV15-22) were treated with either 0.5 μM dimAβ AβO or 1 μM Aβ40 monomers diluted in conditioned neuronal maintenance media for 3 and 24 hours under growth conditions. In addition, cells were treated with a vehicle control (PBS containing 50 mM NaOH and 50 mM HCl). Afterwards cells were fixed and stained as described before^69^.

### Primary neuron culture

Primary neurons were isolated and cultured as described before^69^ with slight modifications. In brief, brains of FVB/N mouse embryos were dissected at embryonic day 13.5. Brainstem and meninges were removed and whole cortex was digested with 1x Trypsin (Panbiotech). Neurons were diluted in pre-warmed (37°C) neuronal plating medium (Neurobasal media (Thermofisher Scientific), 1% FBS, 1x Antibiotic-/Antimycotic solution (Thermofisher Scientific), 1x NS21 (Panbiotech)) and cultivated in a humidified incubator at 37°C, 5% CO_2_. Four days after plating, media was doubled with neuronal maintenance media (Neurobasal media (Thermofisher Scientific), 1x Antibiotic-/Antimycotic solution (Thermofisher Scientific), 1x NS21 (Panbiotech)) and cells were treated with 0.5 μg/ml AraC (Sigma).

### Immunofluorescence staining and data analysis

#### Somatodendritic missorting of Tau

To analyze Tau somatodendritic localization, cells were fixed and stained with a polyclonal rabbit anti-Tau (K9JA, Dako A0024) antibody after Aβ treatment as described before^69^. Fluorescence intensities of cell bodies were quantified using ImageJ software^70,71^. Fluorescence intensity values were normalized to vehicle treated control cells after 3 hours of treatment. All experiments were performed four times; 30 cells were analyzed for each condition. Statistical analysis was done by two-way ANOVA with Tukey’s test for multiple comparisons.

#### Cytotoxic effect of dimAβ

To evaluate AβO toxicity, cells were fixed and stained with NucBlue (Thermofisher Scientific) after dimAβ AβO treatment. Shape and density of nuclei were analyzed and counted: Cells were considered dead, when nuclei appeared condensed and smaller, compared to viable cell nuclei. All experiments were conducted for four times; around 300 nuclei were analyzed for each condition. Statistical analysis was done by two-way ANOVA with Tukey’s test for multiple comparisons.

#### Aβ targeting to postsynaptic spines and calcium imaging

To analyze Aβ binding to synapses, neurons were fixed and stained for F-actin with phalloidin (Thermofisher Scientific) and a monoclonal mouse anti-Aβ (clone 4G8, Merck, #MAB1561) antibody. To monitor spontaneous Ca^2+^ oscillations, primary neurons were labelled with Fluo-4 (Thermofisher Scientific) and Pluronic F127 (Merck) for 20 minutes after 24 hours of dimAβ treatment. Time-lapse movies of different fields were recorded for 1 minute each (framerate: 1 second). Fluorescence intensity changes of cell bodies were quantified over time with ImageJ^70,71^ and corrected for background signal. Fluorescence intensities were normalized to minimum values and peaks per minute were counted for each sample. In total 35 cells were analyzed; statistical analysis was done by unpaired t-test.

### Preparation of fluorescently labeled Aβ for cell culture experiments

#### Preparation of AbberiorStar 520SXP-labeled Cys0-dimAβ

A mutant of dimAβ with an N-Terminal cysteine residue was expressed as described above. For fluorophore-labeling, TCEP-reduced Cys0-dimAβ lyophilisate was incubated in 200mM HEPES pH 7.0 with a two-fold molar excess of maleimide-conjugated AbberiorStar 520SXP fluorophore, which was dissolved in DMF. After 2h of incubation the labeled dimAβ was purified using reverse-phase HPLC. Samples were lyophilized, redissolved in HFIP and aliquots were prepared. These aliquots were lyophilized and stored at RT for later use.

#### Preparation of Aβ42-HiLyte Fluor 647 for cell culture experiments

Aβ42-HiLyte Fluor 647 (Anaspec) was dissolved in HFIP and lyophilized into smaller aliquots (30 μg). For cell culture experiments aliquots were first dissolved in 3 μl 50 mM NaOH. 544 μl phenol red-free DMEM supplemented with 100 U/ml penicillin-streptomycin was added and the pH was recalibrated by the addition of 3 μl 50 mM HCl. To avoid exposure of the Aβ peptide to local low pH environments, the HCl was pipetted into the lid of the tube, closed and quickly vortexed. This procedure yields a 10 μM mostly monomeric stock solution of Aβ42-HiLyte Fluor 647 suitable for cell culture experiments.

#### Preparation of AbberiorStar 520SXP-dimAβ AβOs for cell culture experiments

Abberior STAR 520SXP-labeled AβOs were prepared by dissolving a 1:10 molar ratio of Abberior STAR 520SXP-labeled dimAβ and unlabeled dimAβ in 10 μl 50 mM NaOH. Quickly, 490 μl phenol red-free DMEM supplemented with 100 U/ml penicillin-streptomycin was added and the pH readjusted by adding 10 μl 50 mM HCl. The final dimAβ concentration was 10 μM. The sample was quiescently incubated at 37°C in the dark for 24 h. AβO formation was confirmed using AFM. We were unable to produce large amounts of Abberior STAR 520SXP-labeled dimAβ, therefore unlabeled dimAβ was spiked with labeled dimAβ to achieve concentrations above the COC.

### Neuroblastoma cell culture

SH-SY5Y cells were grown to 80% confluency in DMEM with phenol red, 10 % FBS, and 100 U/ml penicillin-streptomycin in T75 flasks. Experiments were performed in Ibidi collagen IV coated μ-Slide VI 0.4. 7,500 cells (250,000 cells/ml) were seeded into each channel of the slide. Cells adhered to the surface of the channels within an hour of incubation at 37°C, 5% CO_2_, in a humidified chamber. Subsequently, the feeding reservoirs of the channels were filled with further medium. Each day, the medium in the reservoirs was replaced with fresh medium until the cell density was satisfactory for coincubation experiments.

### Coincubation experiments and imaging

For coincubation and imaging experiments phenol red was removed by flushing the channels three times with phenol red-free DMEM supplemented with 100 U/ml penicillin-streptomycin. Subsequently, channels were filled with medium containing corresponding Aβ species. Cells were incubated for 24 hours. Channels were flushed with fresh medium and supplemented with 50 nM Yellow HCK-123 LysoTracker. Imaging was performed either on a Leica Infinity TIRF microscope or on a confocal microscope. Confocal measurements were performed using a TCS SP8 STED 3× (Leica Microsystems) equipped with an HC PL APO CS2 100× objective (NA 1.4) at a scan speed of 600 Hz and a line accumulation of 6. 488 nm of a pulsed white light laser was chosen as excitation for Yellow HCK-123 LysoTracker and AbberiorSTAR520XPS. The emitted fluorescent signal was detected by counting-mode hybrid detectors in the spectral range of 500 – 531 nm for Yellow HCK-123 LysoTracker and 650 – 765 nm for AbberiorStar520SXP. Additionally, a time-gating of 0.1 ns was used to avoid laser reflection.

## Acknowledgements

This project has received funding from the European Research Council under the European Union’s Horizon 2020 research and innovation program, grant agreement No. 726368. We acknowledge support from the Hans und Ilse Breuer-Stiftung, the Else-Kröner-Fresenius Stiftung, and Köln Fortune. We acknowledge the Center of Advanced Imaging (CAI) at the Heinrich Heine University Düsseldorf for providing access to the TCS SP8 STED 3× and support during image acquisition.

## Competing interests

The authors declare no competing interests.

## Notes

### Competing Interest Statement

The authors have declared no competing interest.

