## Supplementary Information for "Endo-lysosomal Aβ concentration and pH enable formation of Aβ oligomers that potently induce Tau missorting"

### Supplementary Figures

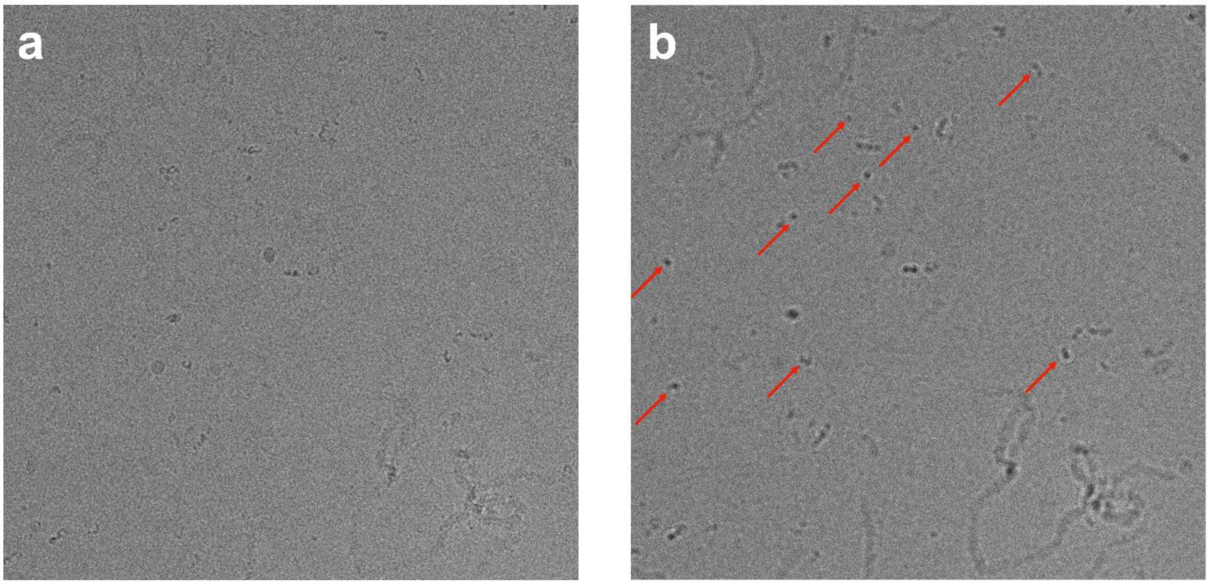


**Supplementary Fig. 1** Representative cryo-EM micrographs of dimAβ AβO. Micrographs were recorded at a defocus of **a** -1.6 µm or **b** -6 µm, respectively. The smallest AβO particles, indicated by red arrows, were selected on the high defocus micrographs for density reconstruction.

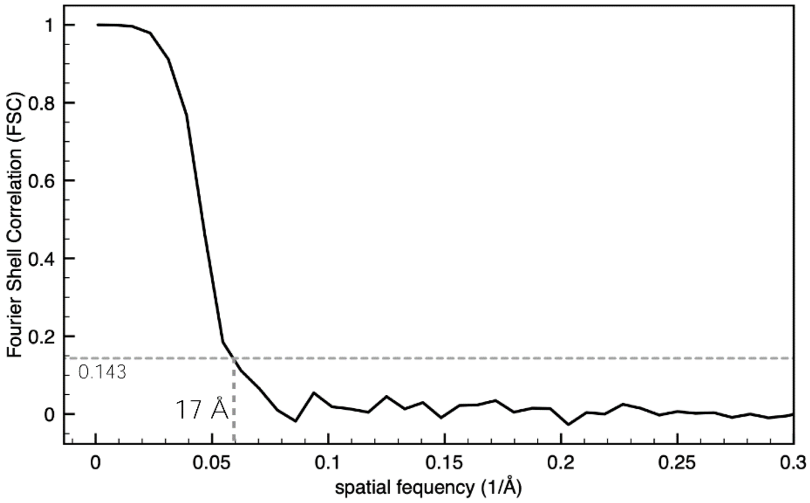


**Supplementary Fig. 2** Fourier shell correlation (FSC) for the 3D reconstruction of the smallest dimAβ AβOs observed on the cryo-EM micrographs yields a resolution estimate of 17 Å.

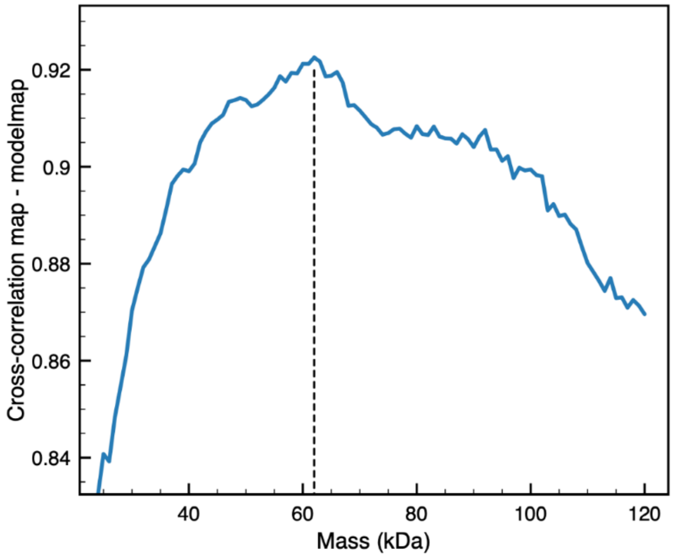


**Supplementary Fig. 3** Density cross-correlation computed for each of the 110 pseudo-atomic model maps with the EM reconstruction after sharpening with VISDEM using the corresponding mass of the pseudo-atomic model. The highest correlation (0.923) is obtained for a mass of 62 kDa.

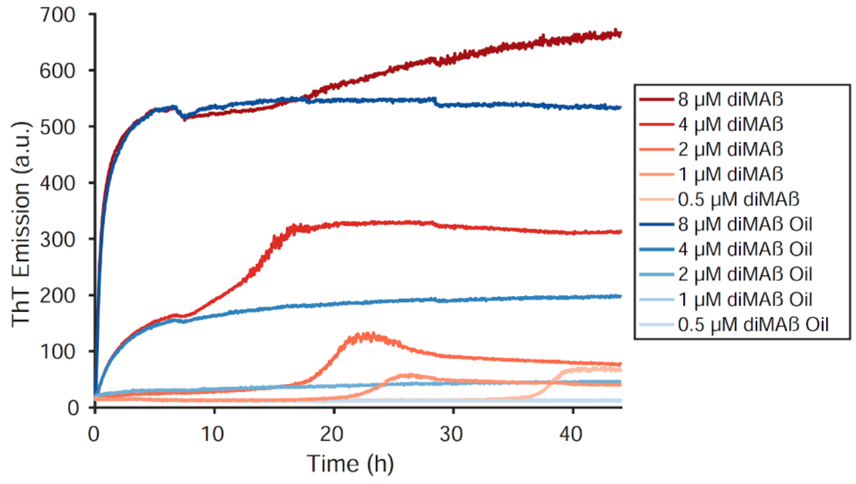


**Supplementary Fig. 4** The air-water interface is crucial for fibril nucleation *in vitro*. Time courses of dimAβ assembly at pH 7.4 monitored by ThT fluorescence in a platereader. Half of the samples were covered by layering 10 µl of mineral oil on top of the aqueous solution. AβO formation was not impaired by mineral oil. Fibril nucleation, on the other hand, was retarded and not detectable during the whole timespan of the experiment.

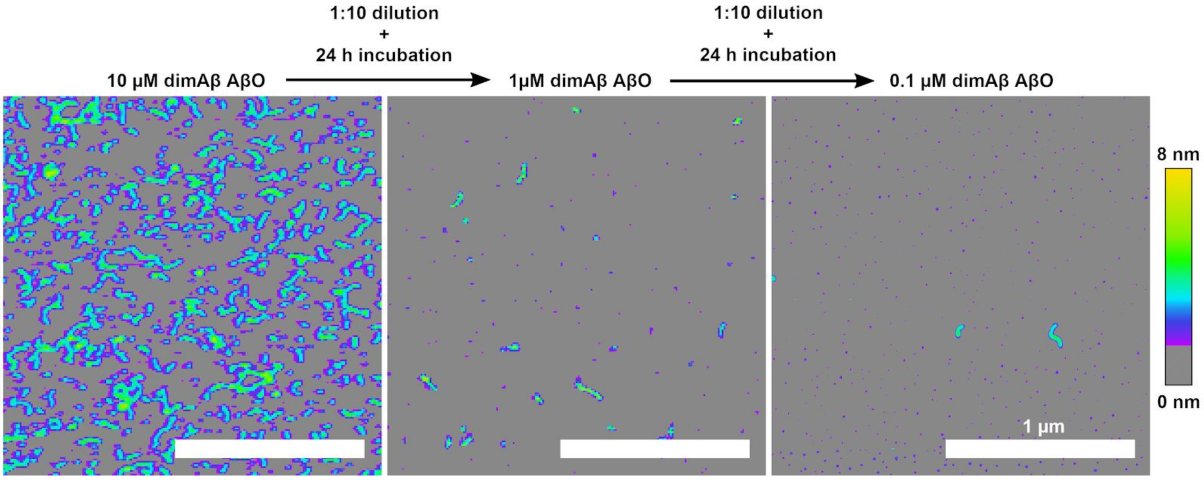


**Supplementary Fig. 5** Long-term stability of diluted AβO. 10 µM dimAβ were quiescently incubated at 37°C, pH 7.4, for 72 h (left). Subsequently, the solution was diluted ten-fold to a dimAβ concentration of 1 µM and further quiescently incubated for 24 h at 37°C (middle). This solution was then further diluted ten-fold to a dimAβ concentration of 0.1 µM, which is far below the COC, and further incubated quiescently for 24 h at 37°C (right). Scalebar, 1 µm.

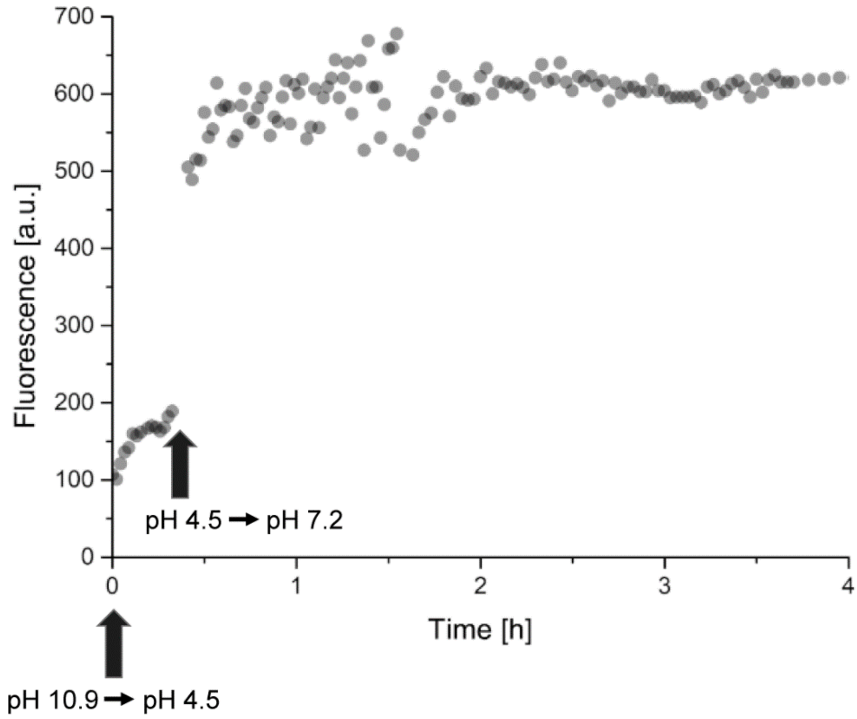


**Supplementary Fig. 6** Stability of AβOs formed from Aβ42 at endo-lysosomal pH after shifting to neutral pH. ThT time course of AβO formation, initiated by pH adjustment from 10.9 to 4.5. Upon pH adjustment from pH 4.5 to pH 7.2 an immediate increase in fluorescence intensity is observed due to the pH sensitivity of ThT fluorescence. Apart from that, no other larger signal changes that would be expected in the case of disassembly of AβOs or replacement of AβOs by an alternative type of aggregate were observed.
